## Supplementary Information for "A method for intuitively extracting macromolecular dynamics from structural disorder"

### Supplementary Material

#### Supplementary Tables

Supplementary Table 1: **Distribution of B-factors for ECHT decompositions of 7k3t, 6xhu, and 6wqf.** The columns contain the average B-factors for each ECHT levels in both Å$^2$ and as a percentage of the total B-factor.

| Level | 7k3t (1.2Å; a-ADPs) Average B (Å$^2$) | 6xhu (1.8Å; TLS + i-ADPs) Average B (Å$^2$) | 6wqf (2.3Å; i-ADPs) Average B (Å$^2$) |
|---|---|---|---|
| Chain | 4.5 (27.9%) | 26.4 (66.3%) | 24.7 (62.7%) |
| Secondary Structure | 4.6 (28.6%) | 4.2 (10.6%) | 9.7 (24.7%) |
| Residue | 3.2 (19.9%) | 3.9 (9.8%) | 3.2 (8.1%) |
| Backbone & Sidechain | 1.6 (10.1%) | 2.7 (6.8%) | 1.1 (2.9%) |
| Atomic | 2.2 (13.5%) | 2.6 (6.5%) | 0.6 (1.5%) |

Supplementary Table 2: **Domain definitions for the spike protein structures 6vxx and 6vyb.** Selections follow the Phenix selection syntax. Domains are defined identically for each chain.

| Group | Selection String |
|---|---|
| 1 | resseq 1:292 |
| 2 | resseq 293:319 or resseq 592:696 |
| 3 | resseq 320:331 or resseq 529:591 |
| 4 | resseq 332:528 |
| 5 | resseq 697:1147 |

Supplementary Table 3: **Distribution of B-factors for ECHT decompositions of 6hcy and 6hd1.** The columns contain the average B-factors for each ECHT levels in both Å$^2$ and as a percentage of the total B-factor.

| Level | 6hcy (3.1Å; group i-ADPs) Average B (Å$^2$) | 6hd1 (3.8Å; group i-ADPs) Average B (Å$^2$) |
|---|---|---|
| Molecule | 44.4 (46.1%) | 40.6 (36.9%) |
| Domain | 31.4 (32.6%) | 38.6 (35.1%) |
| Secondary Structure | 14.5 (15.1%) | 23.3 (21.2%) |
| Residue | 5.3 (5.5%) | 6.8 (6.2%) |
| Atomic | 0.7 (0.7%) | 0.8 (0.7%) |

Supplementary Table 4: **Domain definitions for the STEAP protein structures 6hcy and 6hd1.** Selections follow the Phenix selection syntax. Domains are defined identically for each chain.

| Group | Selection String |
|---|---|
| 1 | resseq 19:195 or resseq 501 |
| 2 | resseq 196:454 or resseq 502:505 |

#### Supplementary Figures

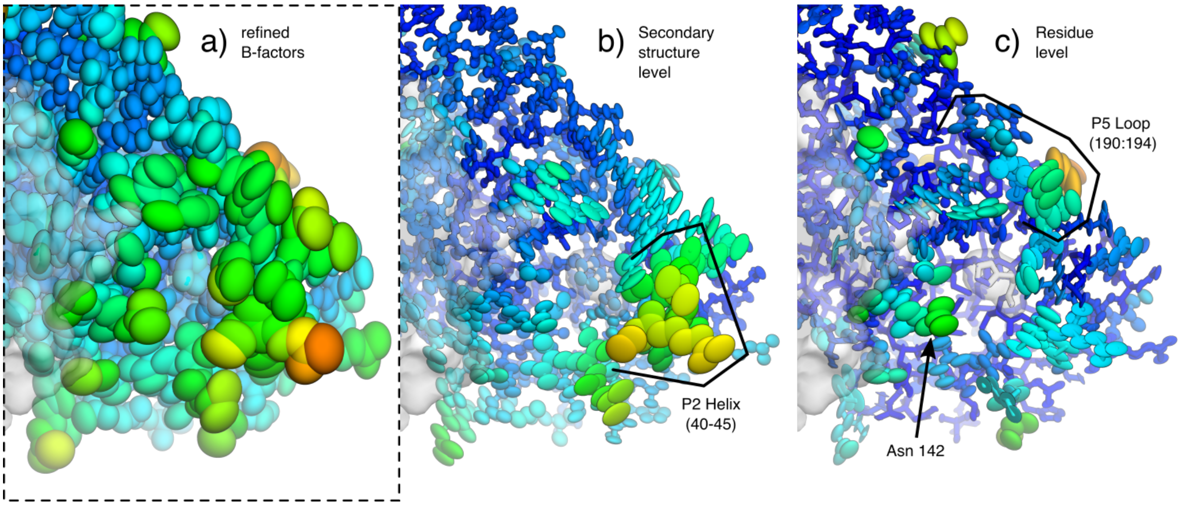


Supplementary Figure 1: **Secondary structure and residue disorder around the catalytic site of the SARS-CoV-2 main protease (7k3t).** Representations and colours are as in Figure 4. (a) Refined B-factors. (b) ECHT secondary structure level. (c) ECHT residue level.

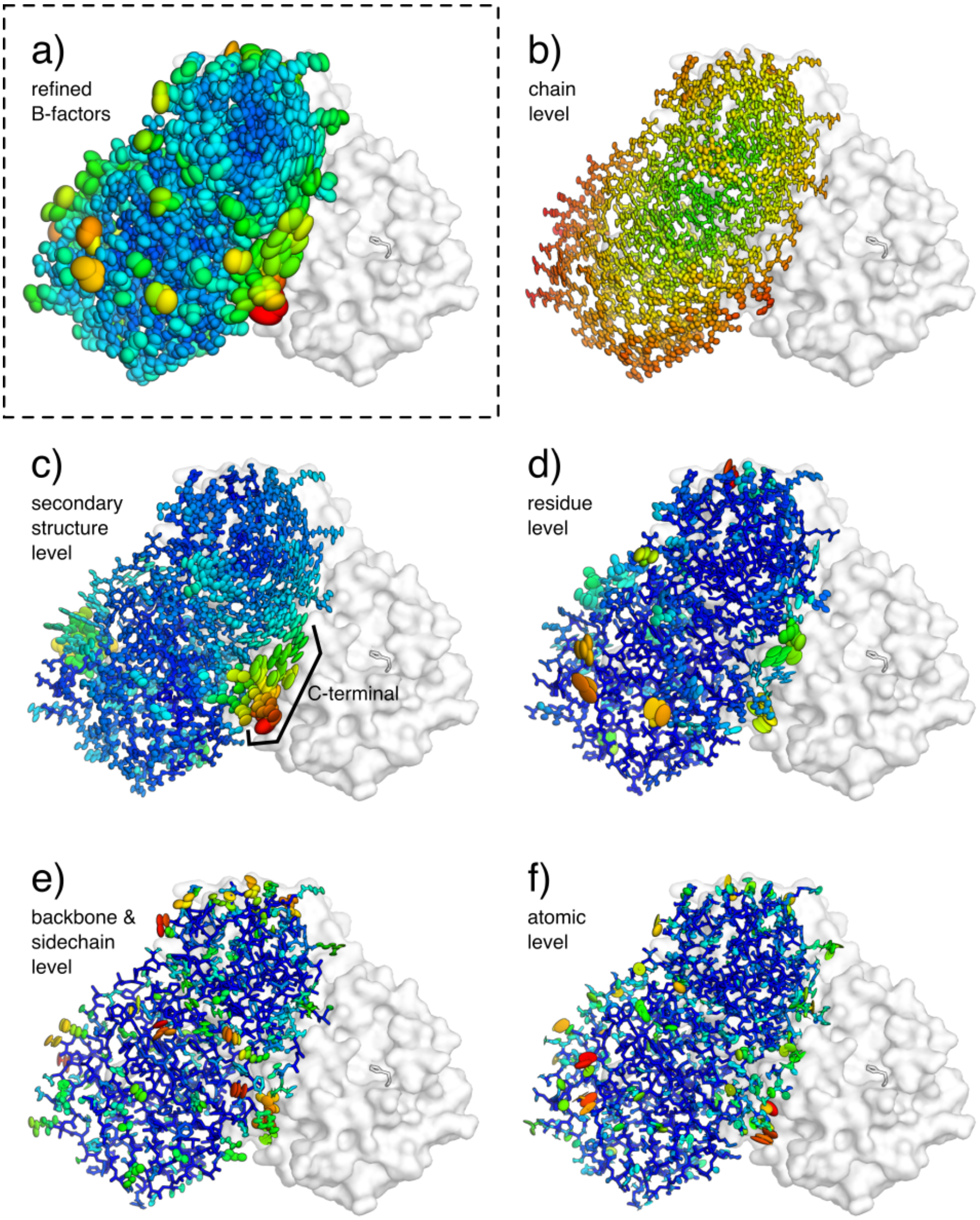


Supplementary Figure 2: **ECHT decomposition of a structure of the SARS-CoV-2 main protease (7k3t; back view).** Representations and colours are as in Figure 4; view is rotated 180° around the y-axis.

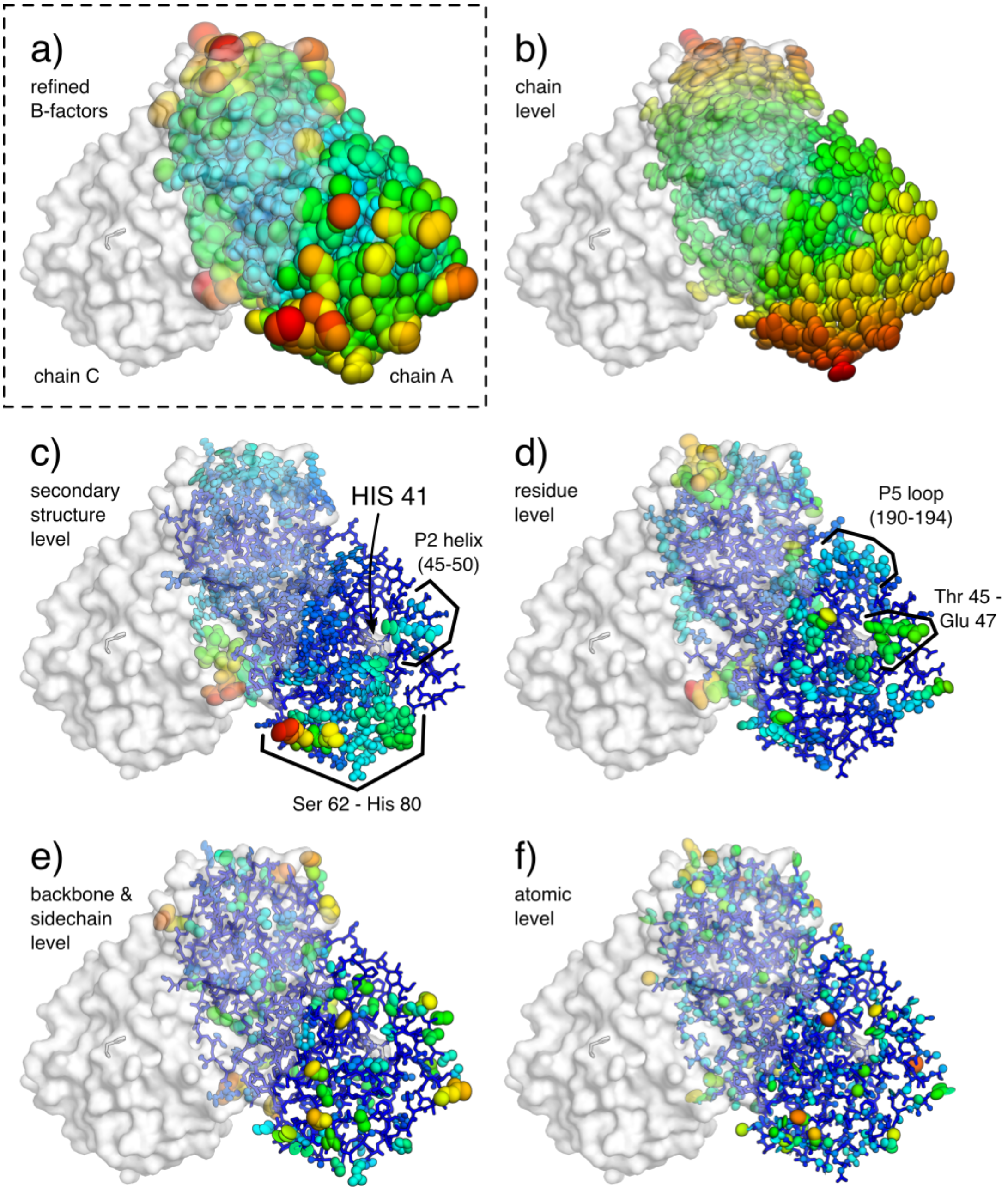


Supplementary Figure 3: **ECHT decomposition of a structure of the SARS-CoV-2 main protease (6xhu).** Chain A is shown as lines and ellipsoids, coloured by B-factor for each structure independently from blue (zero) to green to red (maximum); chain C is shown as semi-transparent surface. The binding site histidine (HIS 41) is also shown as a transparent surface. B-factor ellipsoids are contoured at p=0.95. a) Re-refined structure from PDB_REDO[14] (1.8Å resolution; $R_{work}/R_{free}$ 0.18/0.22; refined with TLS + i-ADPs; B-factors 16.4-128.4Å$^2$). (b-f) Disorder components of each ECHT level (maximum B-factor in brackets): b) chain (58.0Å$^2$), c) secondary structure (36.2Å$^2$), d) residue (40.3Å$^2$), e) backbone & sidechain (40.7Å$^2$) & f) atomic (31.3Å$^2$). (d) Flexible residues line the main binding site. (e) Backbone motions are generally small, with the majority of disorder being isolated to the surface sidechains. (f) Atomic disorder highlights internal motions of residues that cannot be modelled by the rigid body approximation. These principally highlight longer sidechains and backbone carbonyls, as well as presumably absorbing inflated B-factors from modelling errors. All images rendered in pymol[15]

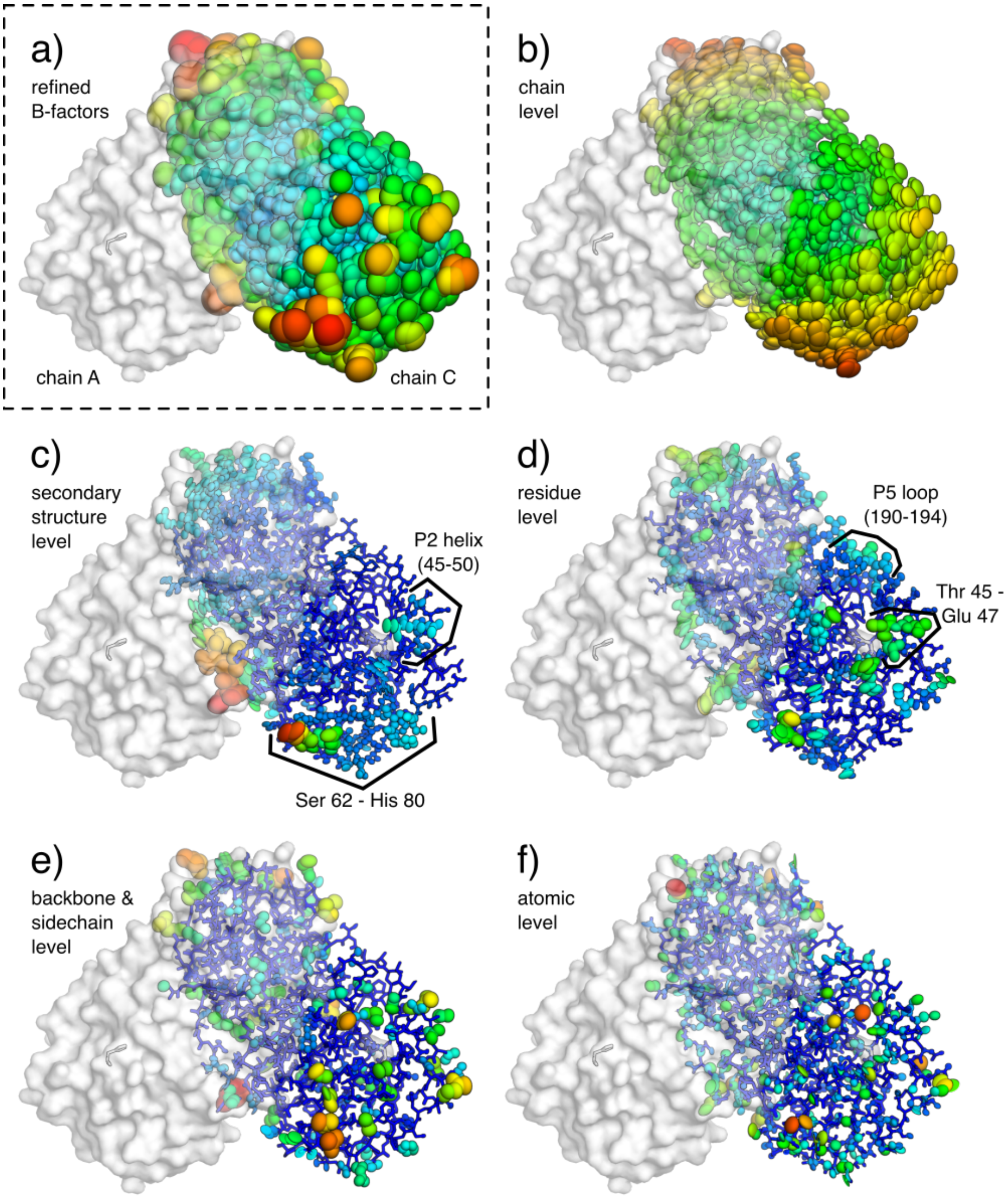


Supplementary Figure 4: **ECHT decomposition of a structure of the SARS-CoV-2 main protease (6xhu, chain C).** Representations and colours are as in Supplementary Figure 3 except for chain C instead of chain A; view is rotated 180° around the y-axis.

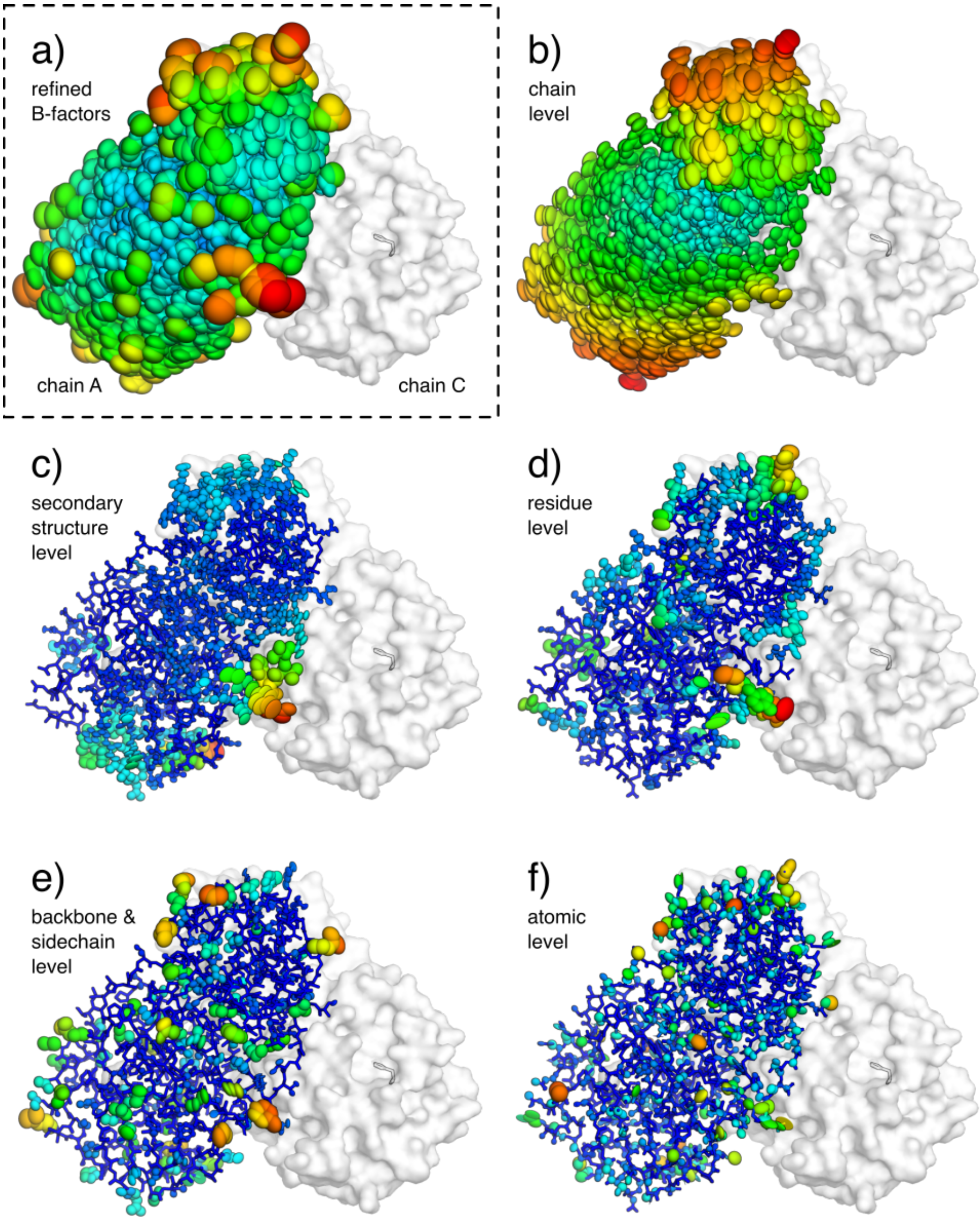


Supplementary Figure 5: **ECHT decomposition of a structure of the SARS-CoV-2 main protease (6xhu, chain A; back view).** Representations and colours are as in Supplementary Figure 3; view is rotated 180° around the y-axis.

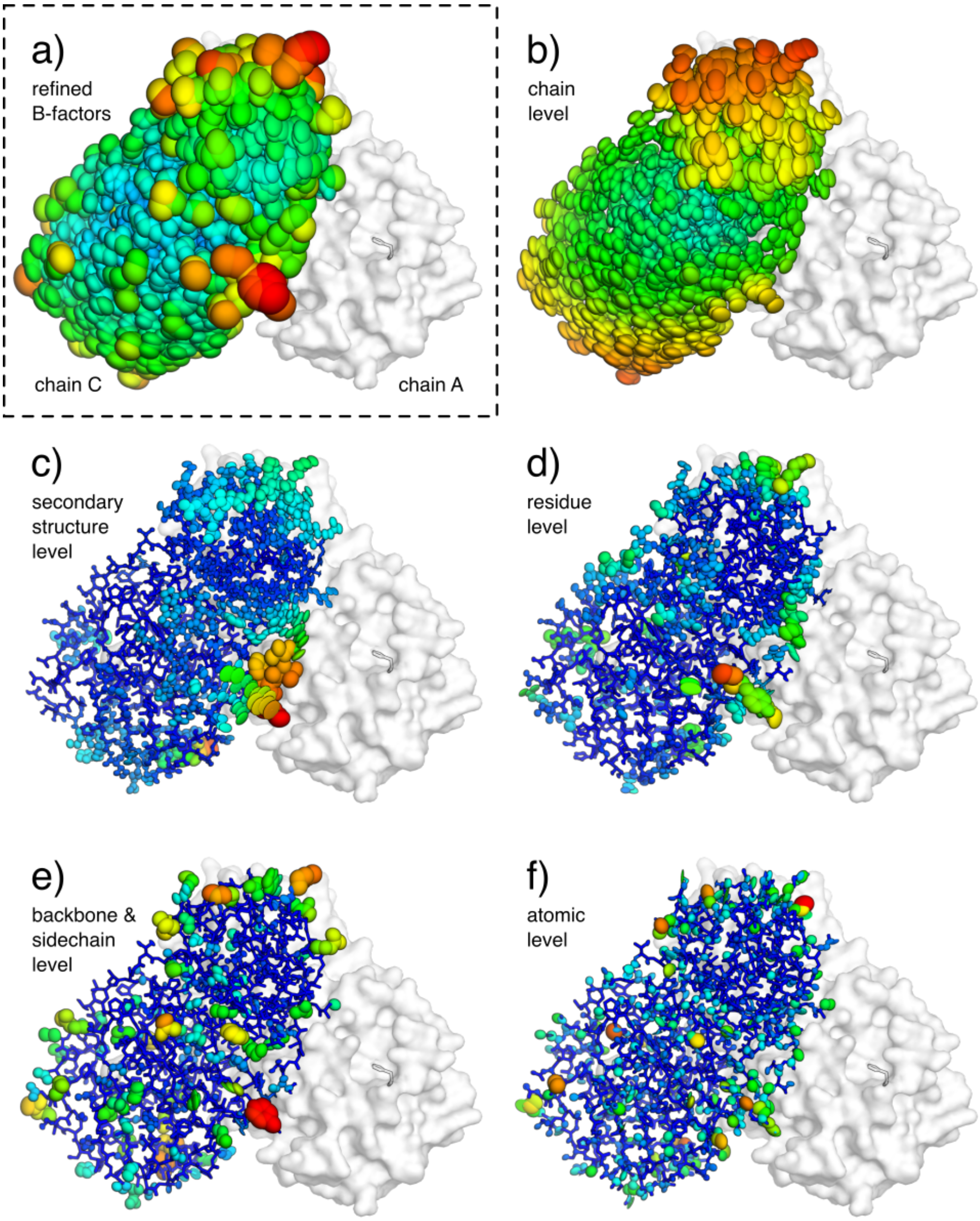


Supplementary Figure 6: **ECHT decomposition of a structure of the SARS-CoV-2 main protease (6xhu, chain C; back view).** Representations and colours are as in Supplementary Figure 3 except for chain C instead of chain A.

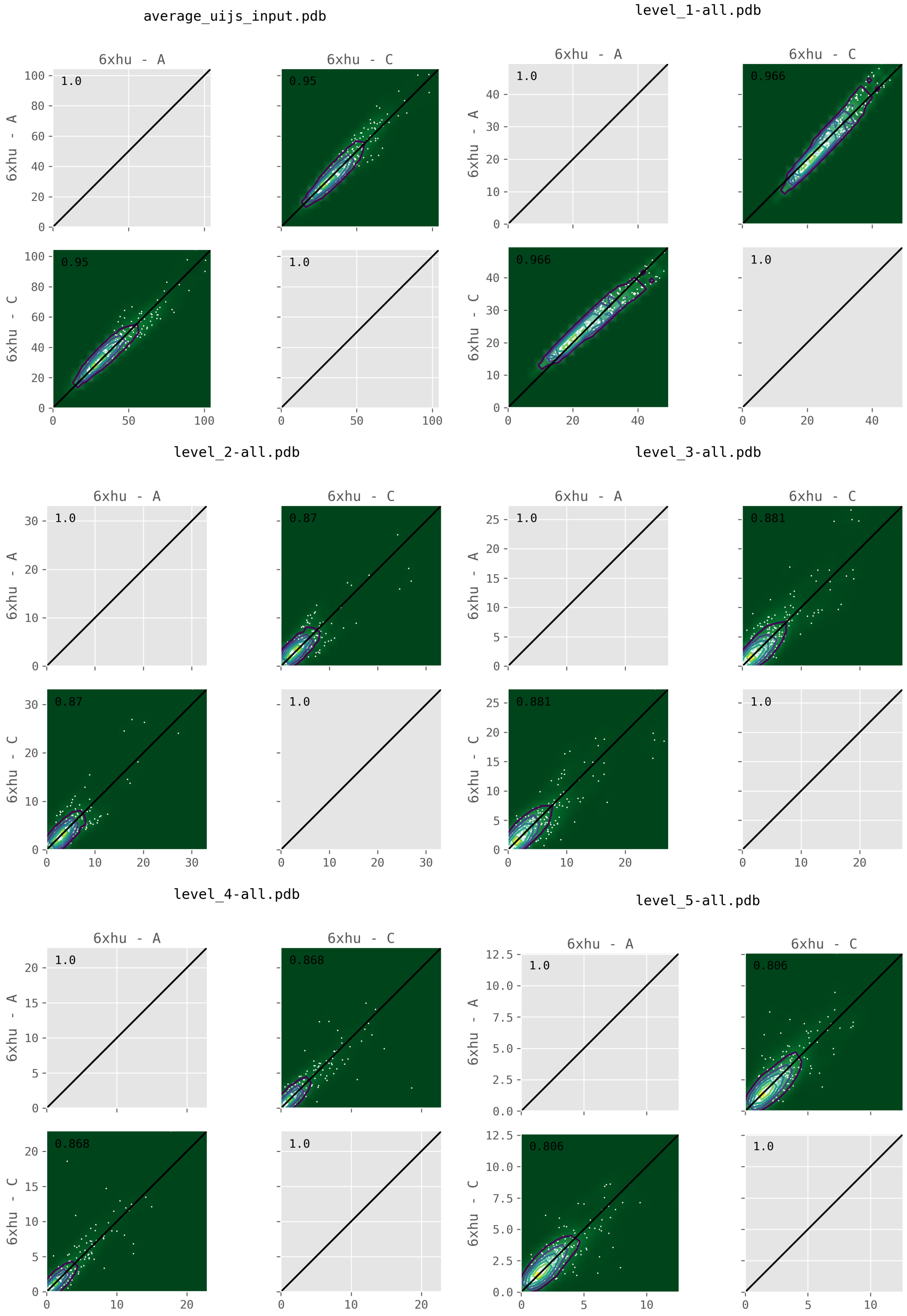


Supplementary Figure 7: **Comparison of B-factors between chains for different levels of ECHT decomposition of 6xhu.** Average B-factors of equivalent residues are compared between chains. Contour plots are also shown to account for over-plotting of small B-factor regions. (a) Total B-factor, (b) chain level, (c) secondary structure level, (d) residue level (e) backbone/sidechain level and (f) atomic level. Inset values: correlation coefficients between the two sets.

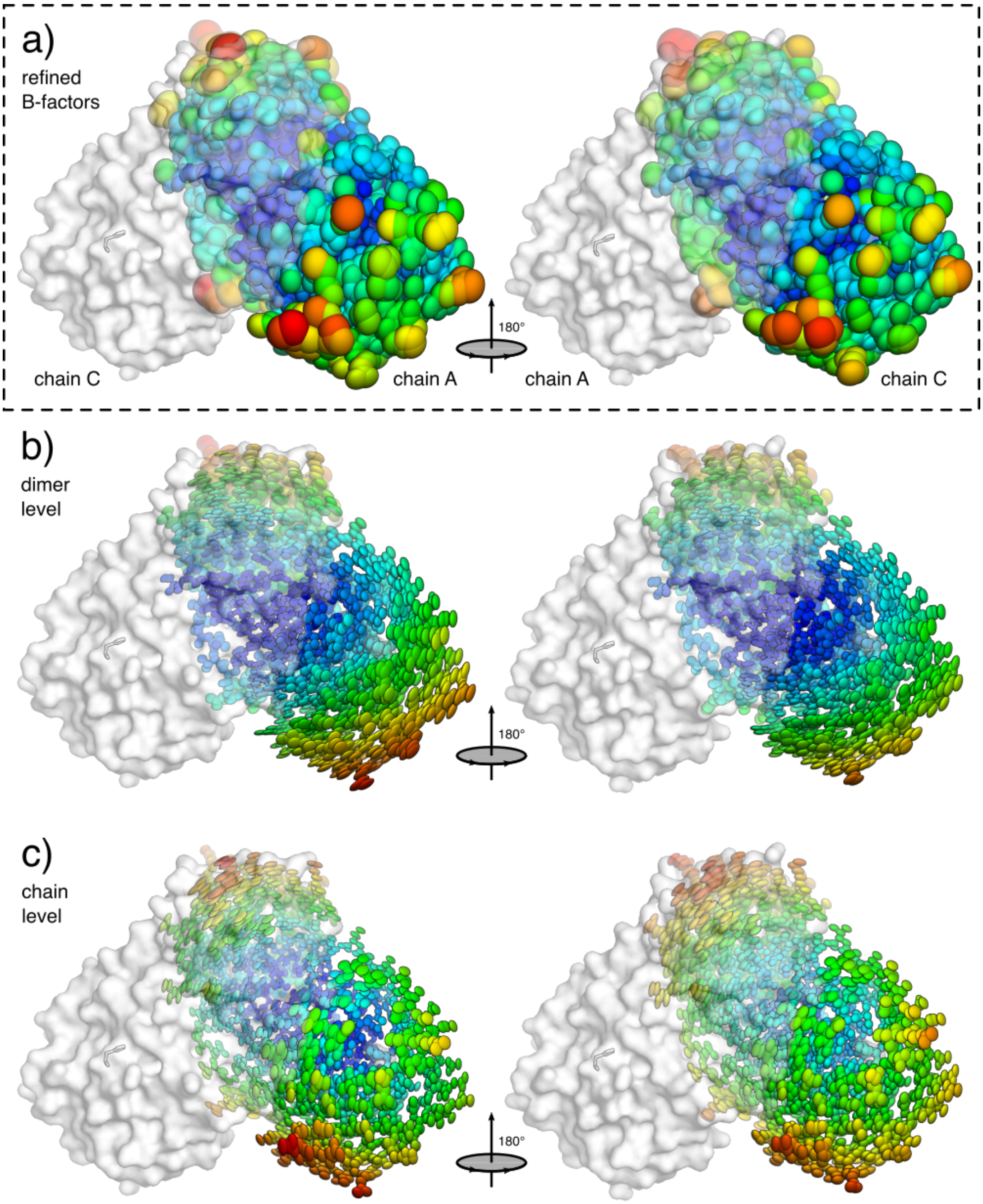


Supplementary Figure 8: **ECHT decomposition of 6xhu with additional dimer level.** Structures are shown coloured by B-factors from blue (minimum) to green to red (maximum). (a) The PDB_REDO re-refined structure (TLS + iADPs; one TLS group per chain; B-factors 16.4-128.4Å$^2$). The TLS components groups are the sum of any dimer disorder and any monomer disorder. An additional level was added to the ECHT description corresponding to the dimer of both chains. (b) The new dimer level results in a mostly-rotational component around the dimer's centre of mass (B-factors 6.8-35.2Å$^2$). (c) The new chain components, after determination of the joint molecular component, now show more clearly a rocking/rolling motion around the extended dimer interface (B-factors 3.1-23.8Å$^2$).

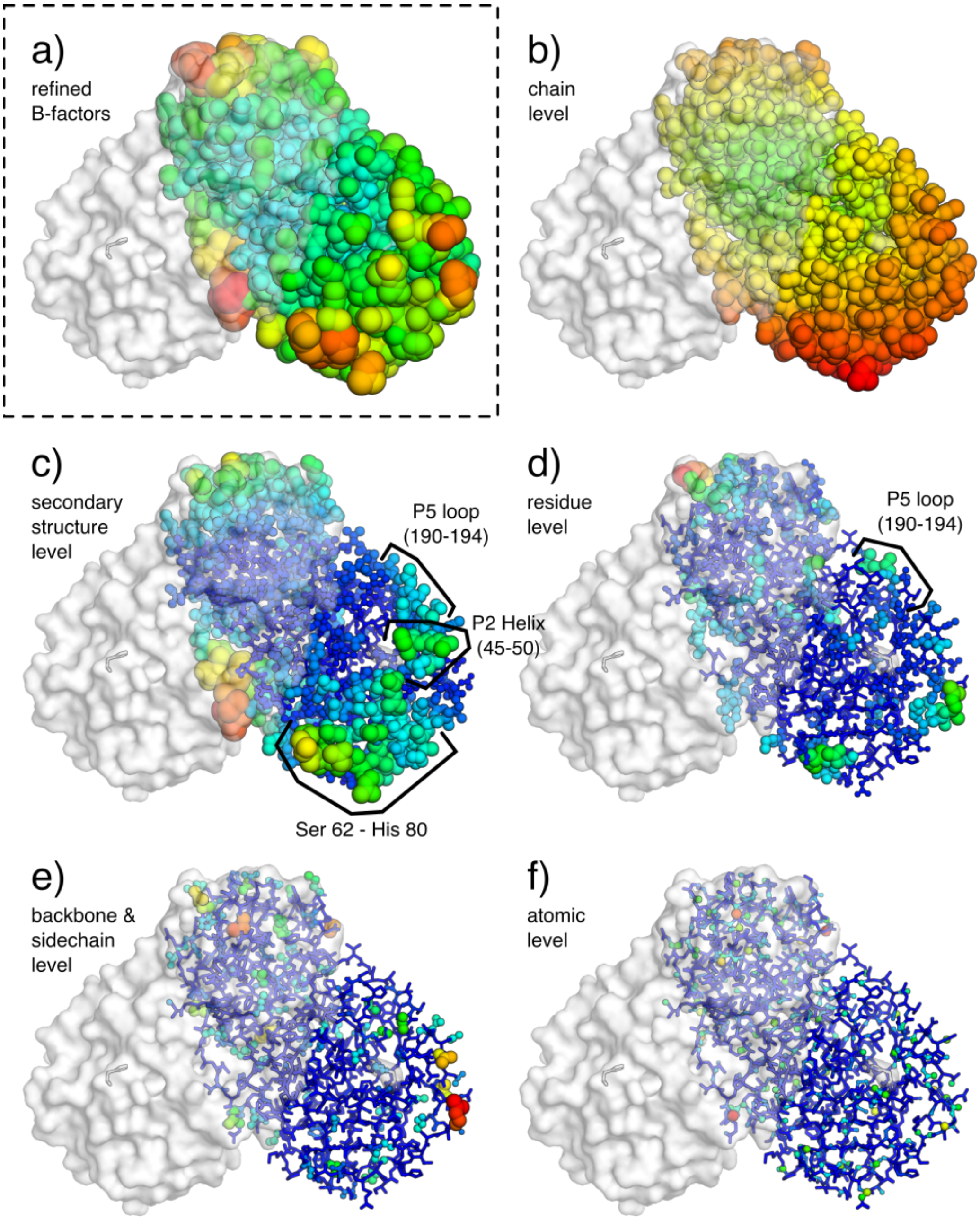


Supplementary Figure 9: **ECHT decomposition of a structure of the SARS-CoV-2 main protease (6wqf).** Layout and representations are as in Figure 4. (a) Re-refined structure from PDB_REDO[14] (2.3Å resolution; $R_{work}/R_{free}$ 0.17/0.23; refined with i-ADPs; B-factors 12.5-108.5Å$^2$). (b-f) Disorder components of each ECHT level (maximum B-factor in brackets): (b) chain (40.5Å$^2$), (c) secondary structure (66.8Å$^2$), (d) residue (46.3Å$^2$), (e) backbone & sidechain (21.8Å$^2$) & (f) atomic (10.6Å$^2$). (b) The chain components once more indicate a rolling/rocking motion around the dimerisation interface (c.f. Supp. Figure reffig:6xhu-array). (c) Residues 62-78 are again identified as flexible at the secondary structure level (c.f. Supp. Figure reffig:6xhu-array), as is the stretch of residues lining the catalytic site. See also Supp. Figure 10 for back view.

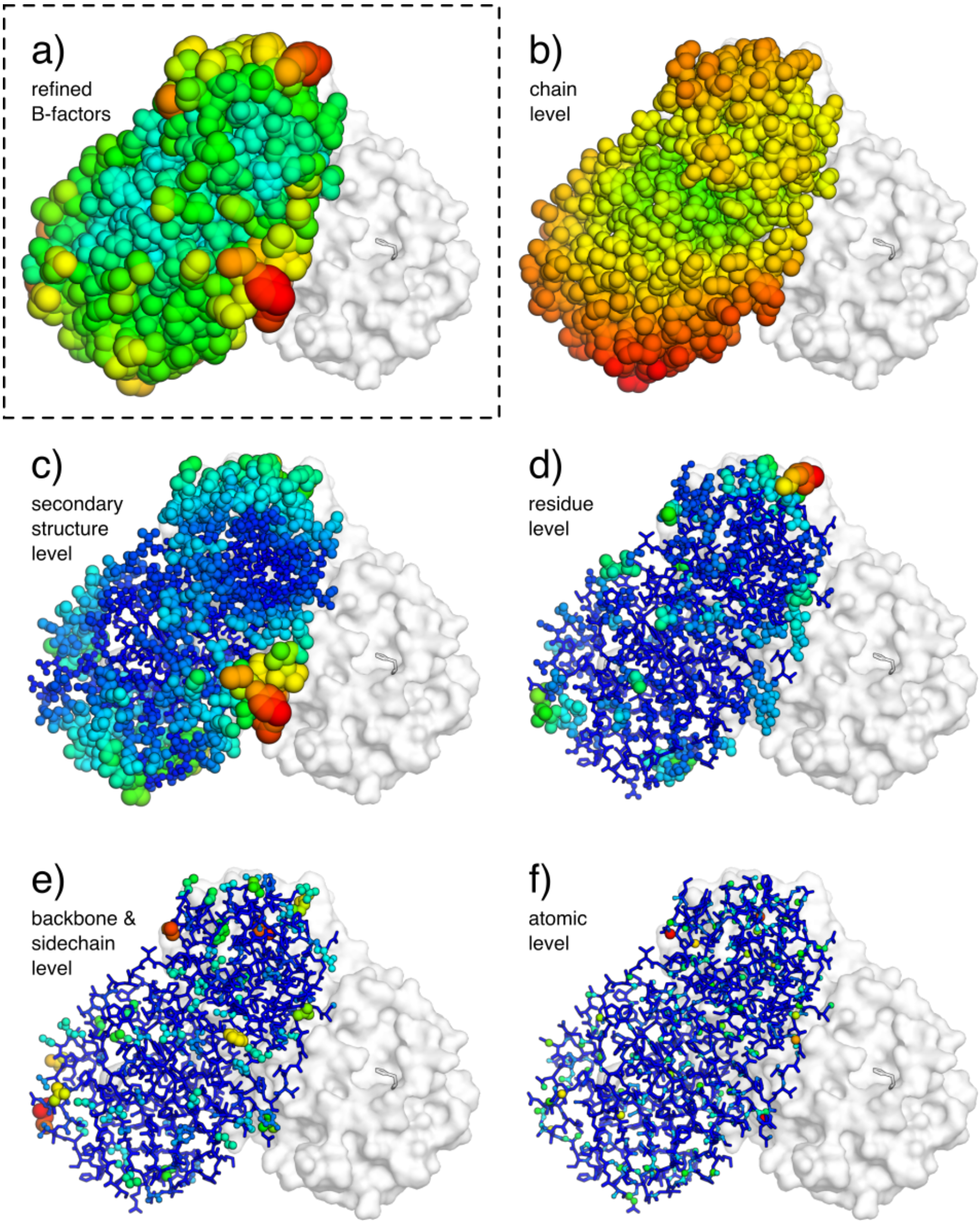


Supplementary Figure 10: **Decomposition of 6wqf (back view).** Representations and colours are as in Supplementary Figure 9; view is rotated 180° around the y-axis.

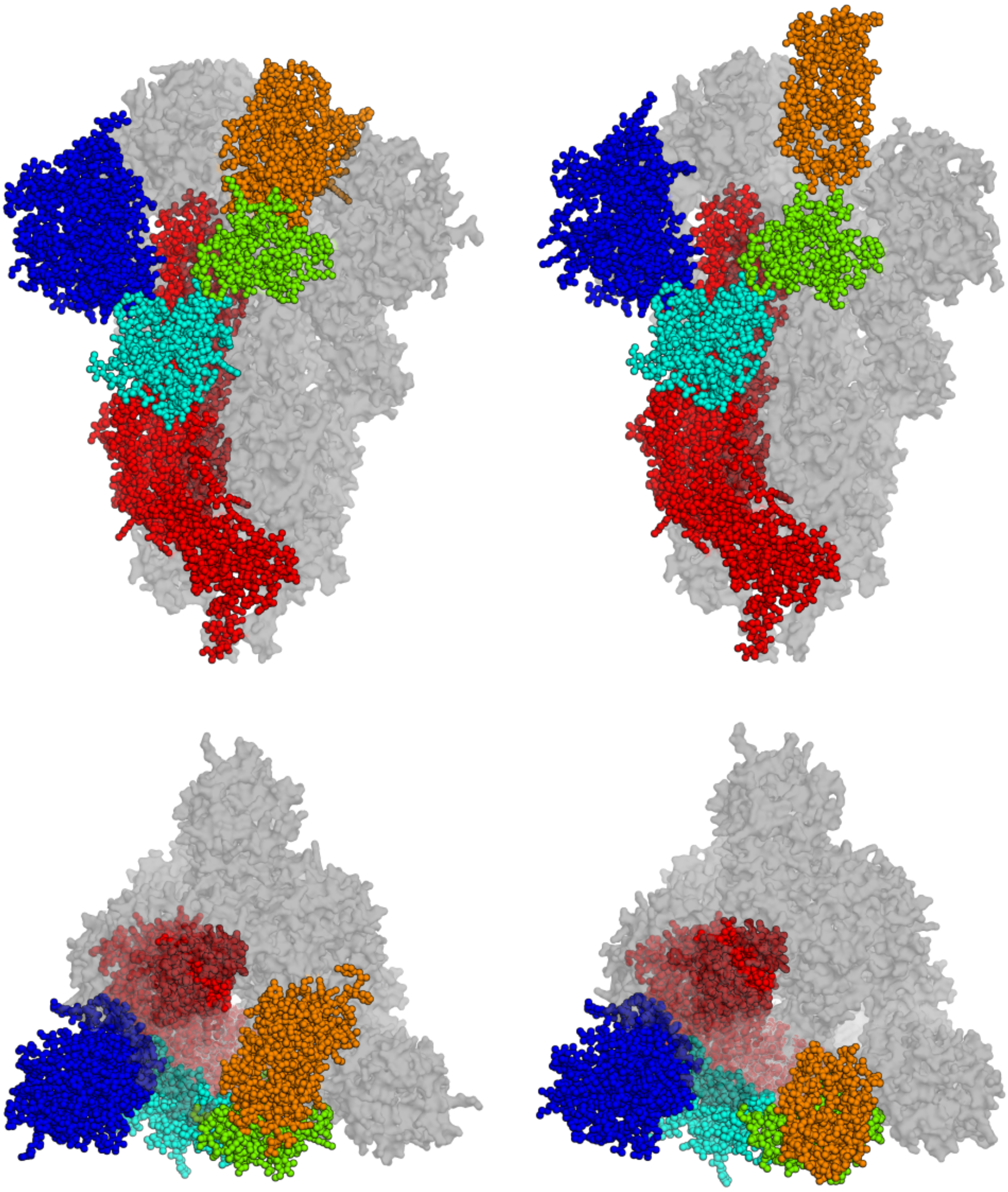

Supplementary Figure 11: **Domain definitions for SARS-CoV-2 surface glycoprotein (spike).** The different domains used in the "domain" level of the ECHT decompositions, as defined in Supp. Table 2, are shown in different colours for one chain of 6vxx (left) and 6vyb (right). Orientations and configuration as in Figure 5.

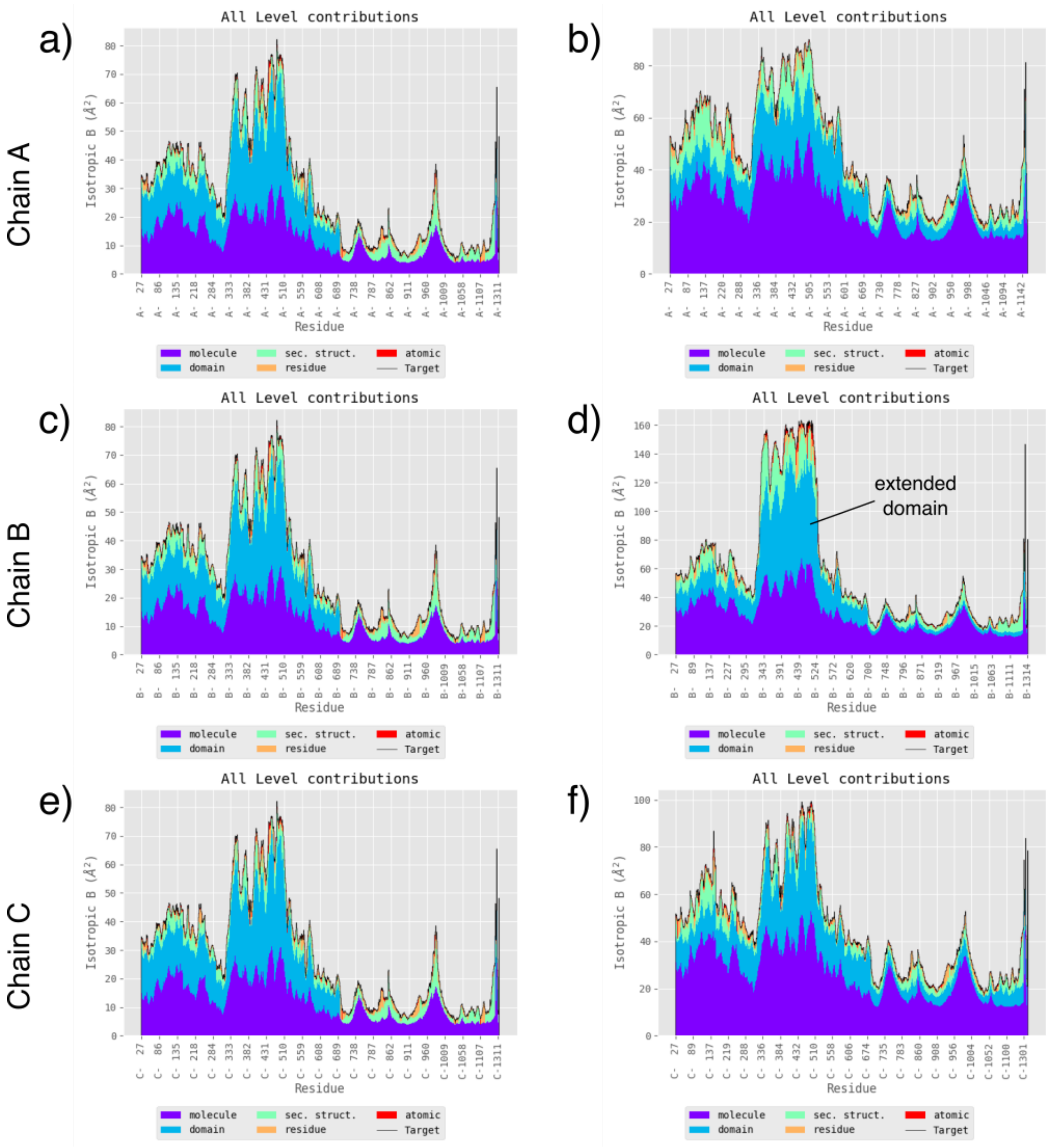


Supplementary Figure 12: **B-factor profiles for ECHT decomposition of two structures of SARS-CoV-2 surface glycoprotein (6vxx and 6vyb).** (a,c,e) Disorder profiles of 6vxx by chain. (b,d,f) Disorder profiles of 6vyb by chain.

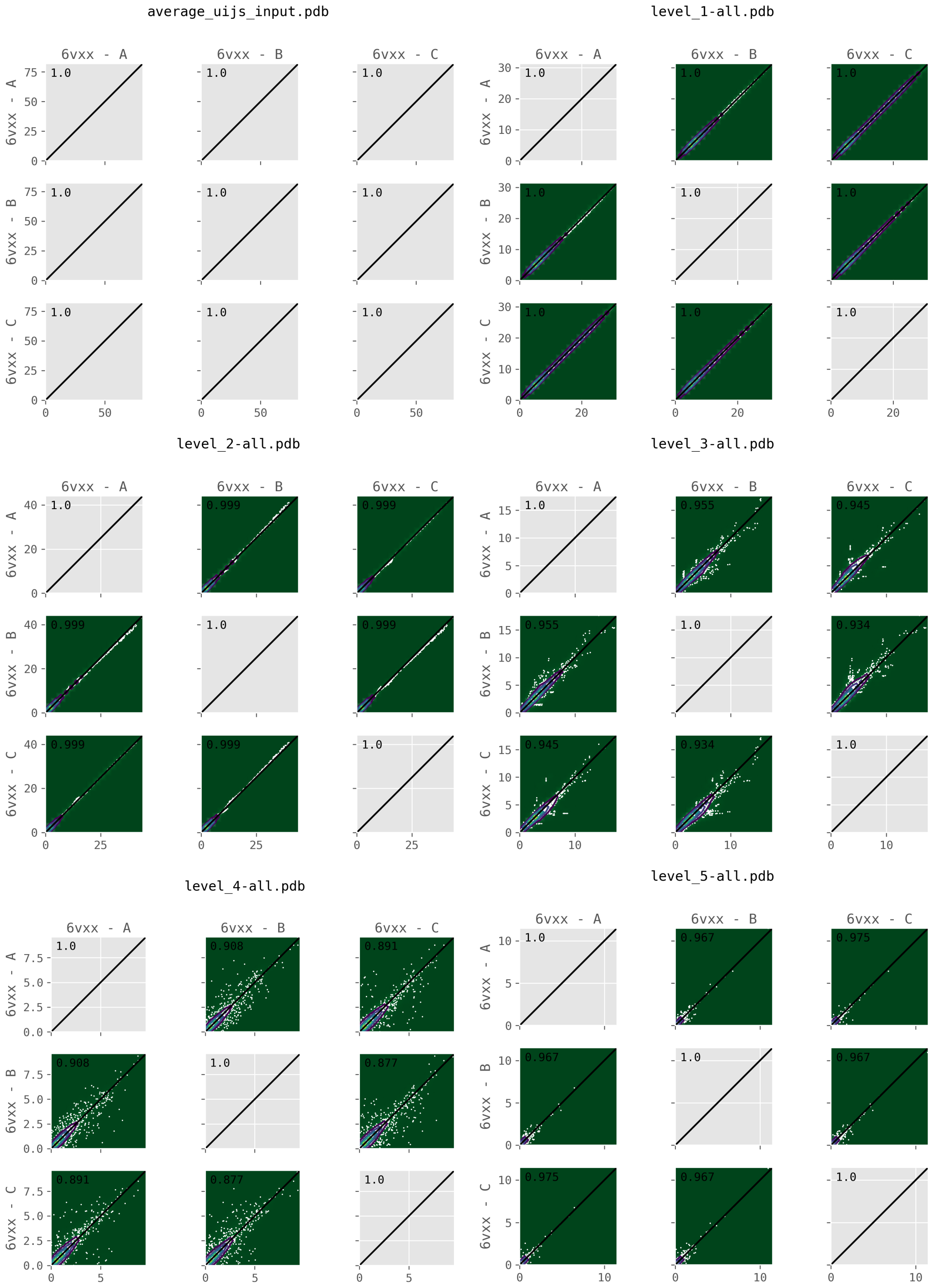


Supplementary Figure 13: **Comparison of atomic B-factors between chains for different levels of ECHT decomposition of 6vxx.** Average B-factors of residues are compared between chains. Contour plots are shown to account for over-plotting of small B-factor regions. (a) Total B-factor, (b) molecule level (c) domain level, (d) secondary structure level, (e) residue level, and (f) atomic level. Inset values: correlation coefficients between the two sets.

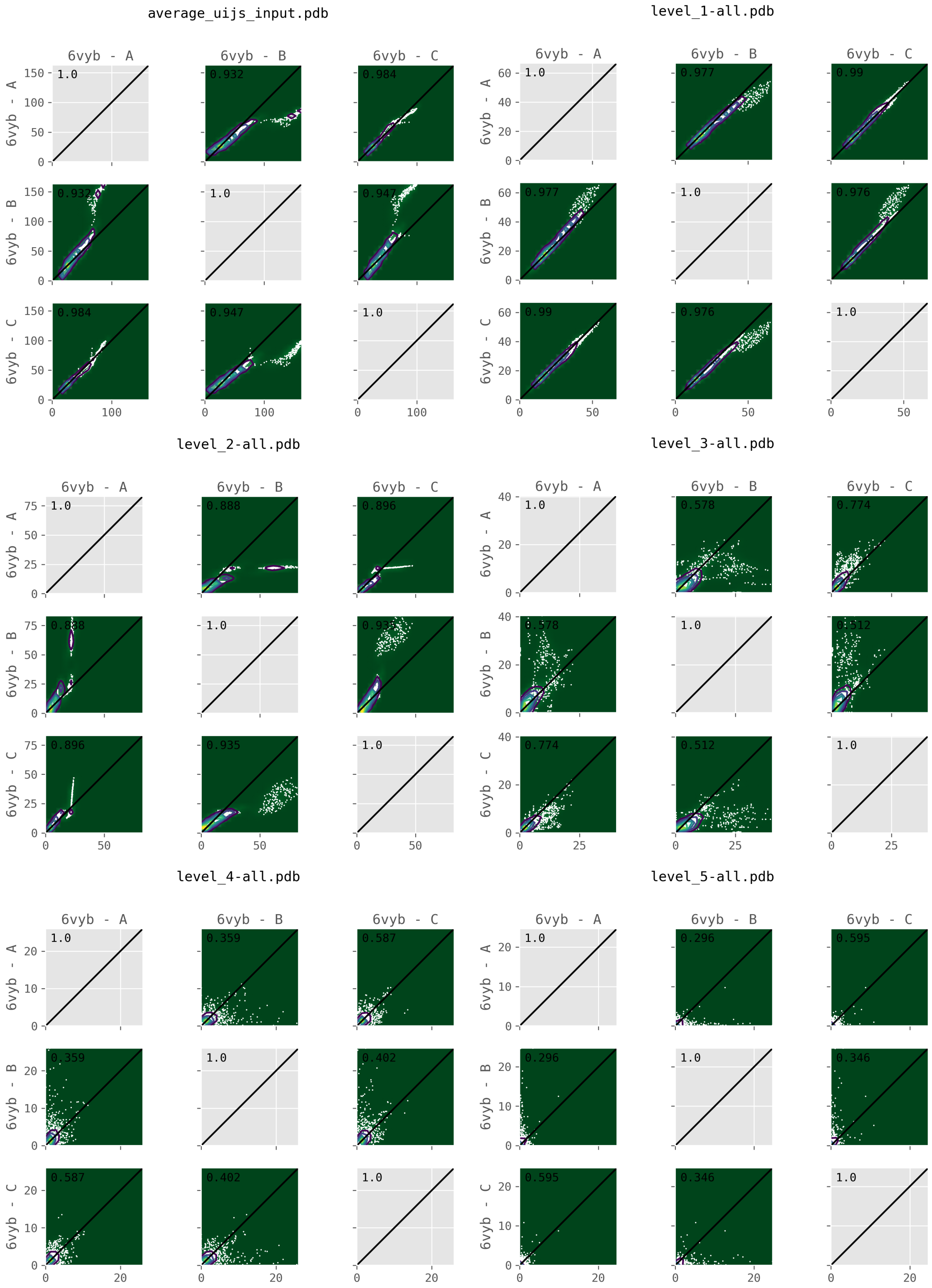


Supplementary Figure 14: **Comparison of atomic B-factors between chains for different levels of ECHT decomposition of 6vyb.** Average B-factors of residues are compared between chains. Contour plots are shown to account for over-plotting of small B-factor regions. (a) Total B-factor, (b) molecule level (c) domain level, (d) secondary structure level, (e) residue level, and (f) atomic level. Inset values: correlation coefficients between the two sets.

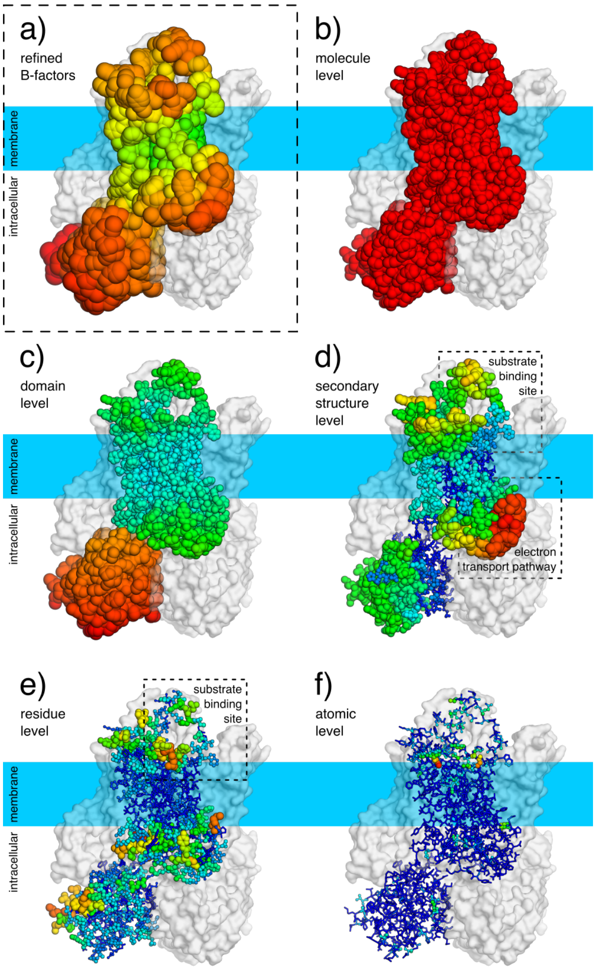


Supplementary Figure 15: **ECHT decomposition of STEAP4 (6hcy; side view).** Representations, layout and colouring as in Figure 6. (a) Deposited structure (3.1Å resolution; one i-ADP per residue; B-factors 55.4-146.7Å$^2$). (b-f) Disorder components of each ECHT level (maximum B-factor in brackets): (b) molecule (44.7Å$^2$), (c) domain (65.8Å$^2$), (d) secondary structure (51.7Å$^2$), (e) residue (28.7Å$^2$), & (f) atomic (13.8Å$^2$).

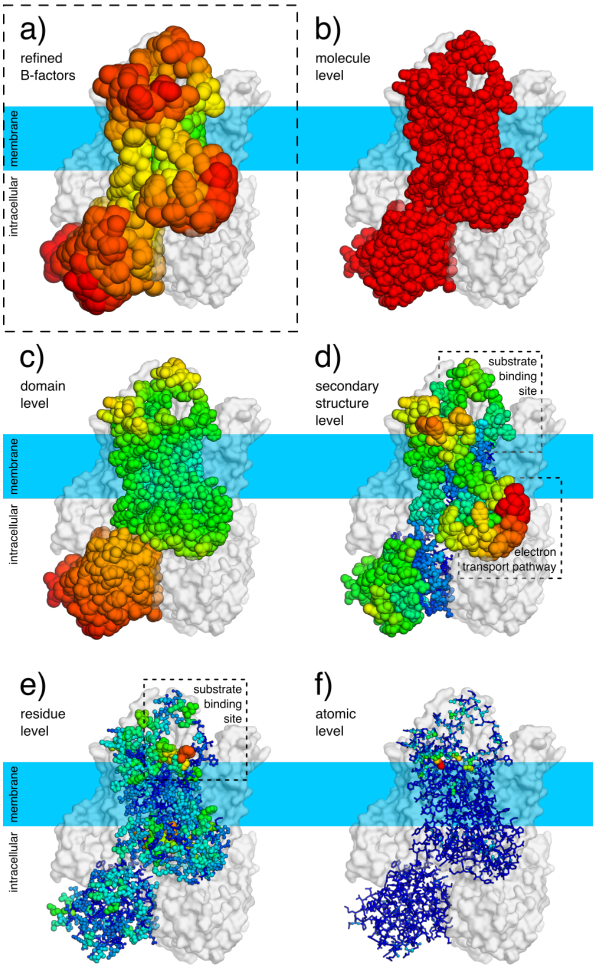


Supplementary Figure 16: **ECHT decomposition of STEAP4 (6hd1; side view).** Representations, layout and colouring as in Figure 6. (a) Deposited structure (3.8Å resolution; one i-ADP per residue; B-factors 55.5-158.8Å$^2$). (b-f) Disorder components of each ECHT level (maximum B-factor in brackets): (b) molecule (42.3Å$^2$), (c) domain (74.0Å$^2$), (d) secondary structure (74.5Å$^2$), (e) residue (41.2Å$^2$), & (f) atomic (18.9Å$^2$).

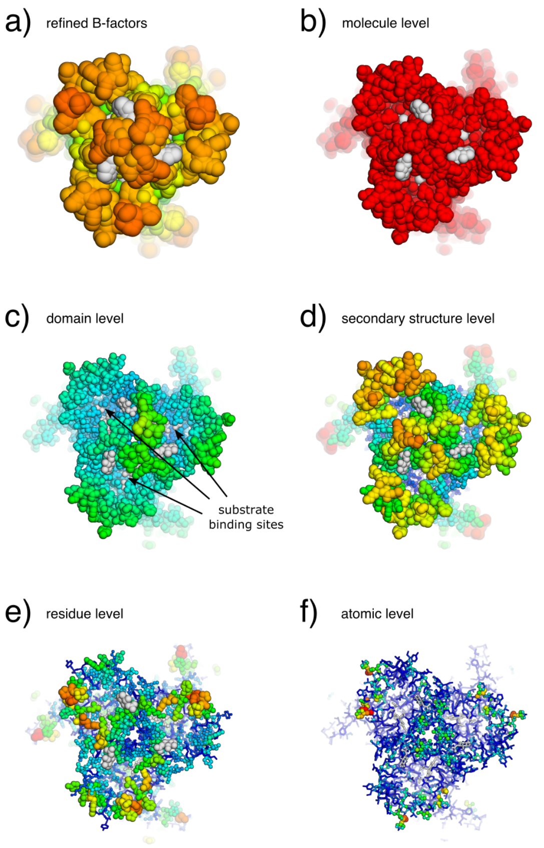


Supplementary Figure 17: **ECHT decomposition of STEAP4 (6hcy; top view).** Representations, layout and colouring as in Supp. Figure 15, except with ellipsoids show for all chains and non-protein atoms coloured grey; view is rotated 90° around the x-axis.

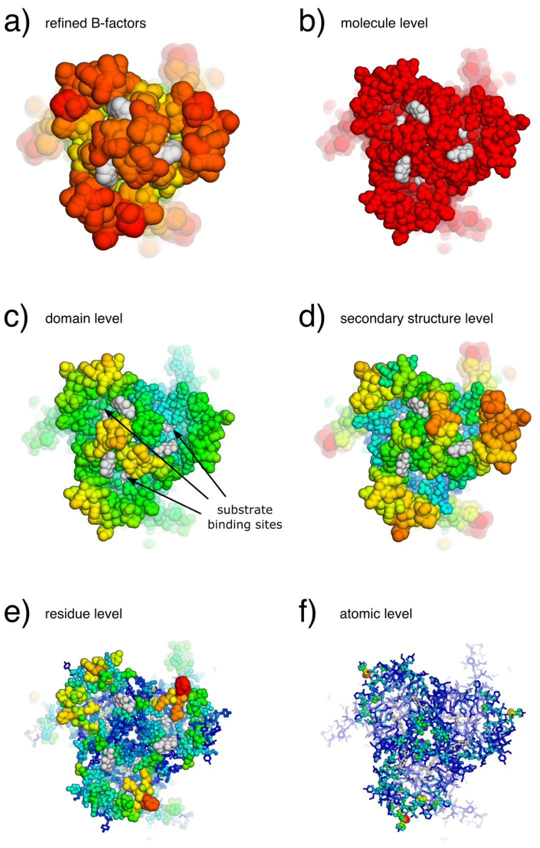


Supplementary Figure 18: **ECHT decomposition of STEAP4 (6hd1; top view).** Representations, layout and colouring as in Supp. Figure 16, except with ellipsoids show for all chains and non-protein atoms coloured grey; view is rotated 90° around the x-axis.

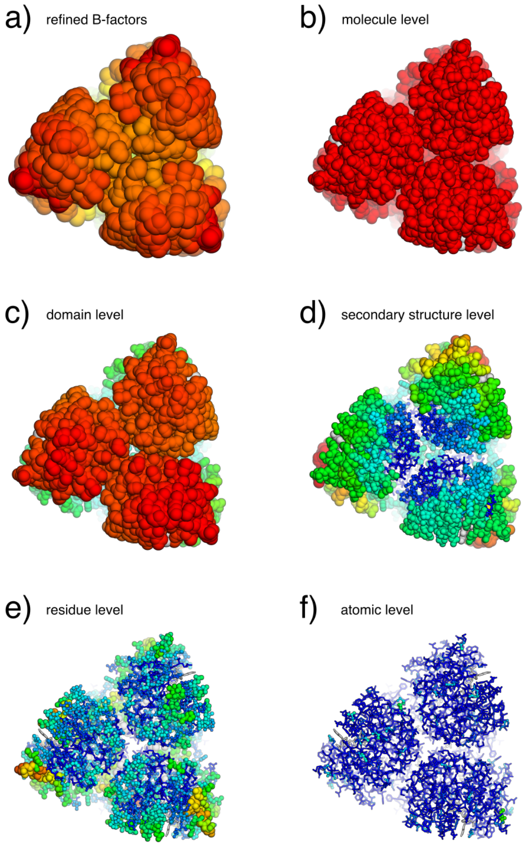


Supplementary Figure 19: **ECHT decomposition of STEAP4 (6hcy; bottom view).** Representations, layout and colouring as in Supp. Figure 15, except with ellipsoids show for all chains; view is rotated -90° around the x-axis. A sharp transition is observed in the disorder pattern in the secondary structure level in the middle of the intracellular domain. NADPH binds to this domain; the flexibility in this domain may permit for the turnover of NADPH molecules.

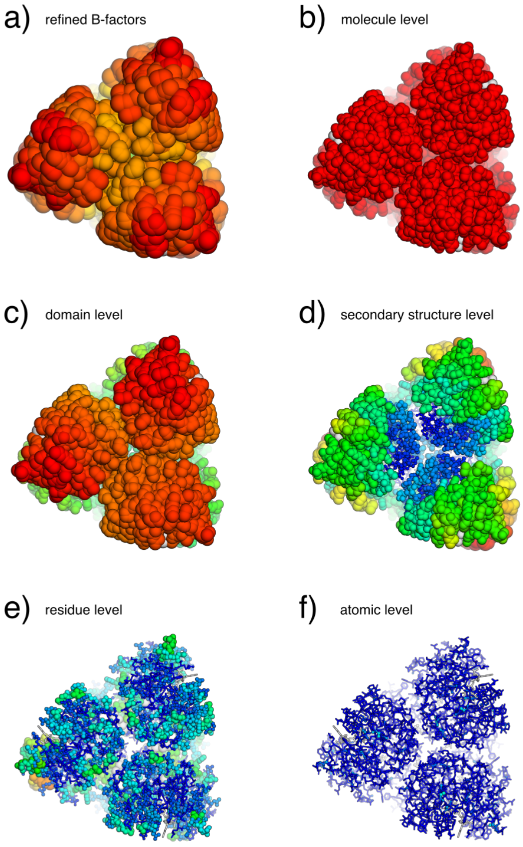


Supplementary Figure 20: **ECHT decomposition of STEAP4 (6hd1; bottom view).** Representations, layout and colouring as in Supp. Figure 16, except with ellipsoids show for all chains; view is rotated -90° around the x-axis. A sharp transition is observed in the disorder pattern in the secondary structure level in the middle of the intracellular domain. NADPH binds to this domain; the flexibility in this domain may permit for the turnover of NADPH molecules.

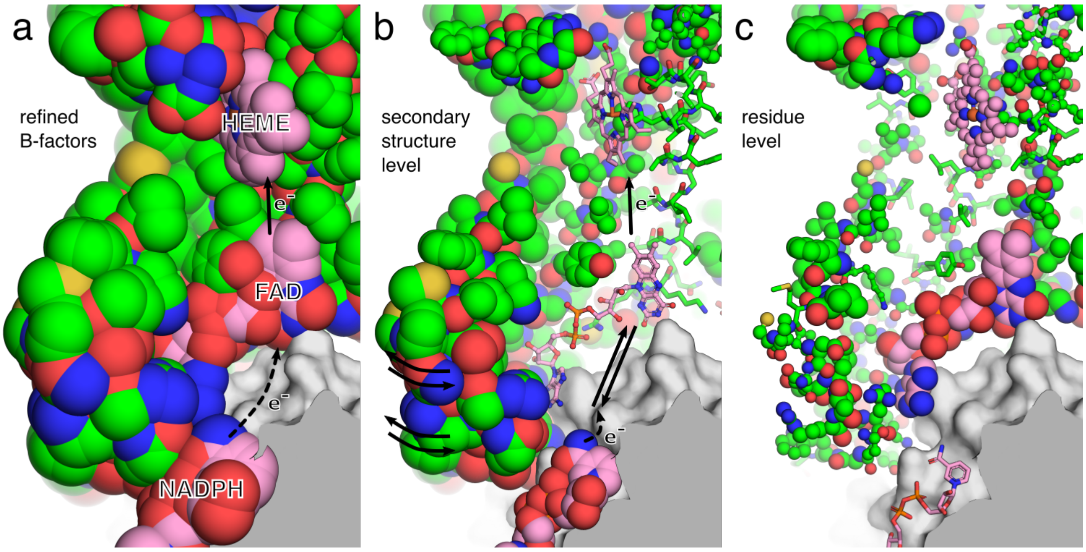


Supplementary Figure 21: **Putative flexibility-based mechanism of electron transport in STEAP4 (6hd1).** Layout and representations as in Figure 7. In the secondary structure level, ligands are assigned to the nearest group. The FAD molecule in 6hd1 is assigned to a different group than in 6hcy, which causes it to pick up a smaller secondary-structure component than in Figure 7. However the same pattern is present: the FAD molecule has excess disorder relative to the surrounding ordered protein.

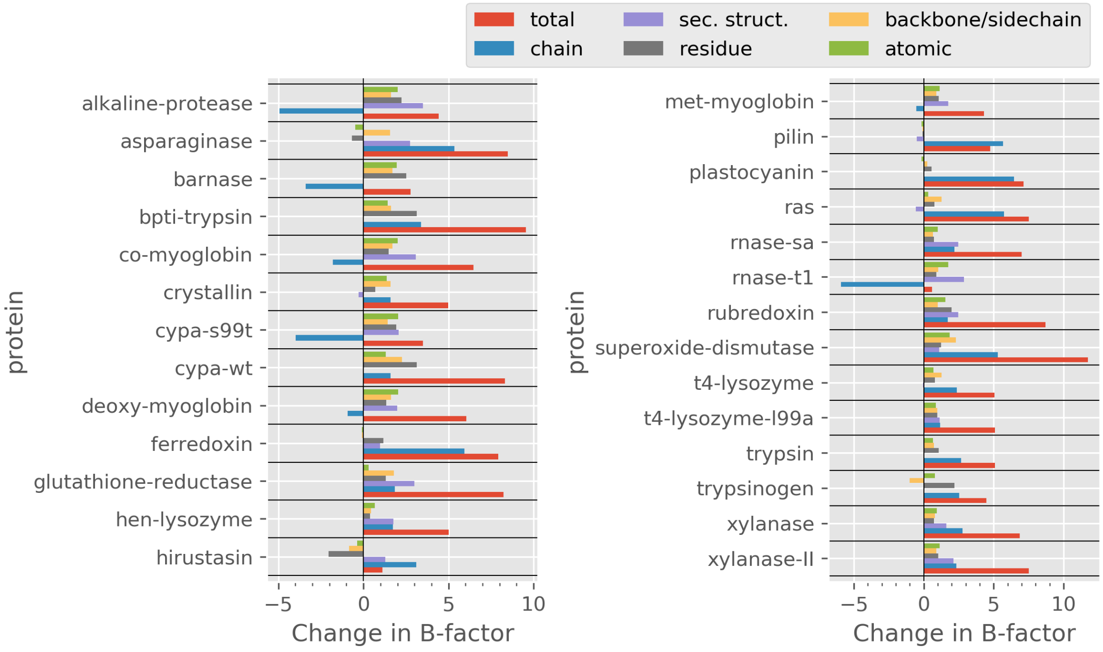


Supplementary Figure 22: **Comparison of ECHT decompositions between structures at different temperatures.** The change in the average B-factors for each level in ECHT decompositions within pairs of structures at cryogenic and room temperatures. The increase in global average B-factor shows a “warming up” of the structure, though this may also be a factor of resolution of the data. The increase in average B-factor of small-scale levels shows an increase of distinct disorder at these levels, and in some case reduces the disorder of the chain level as components that are similar at cryogenic temperatures (thereby becoming present in the lower levels) become distinct at higher temperatures and therefore modelled at higher levels.

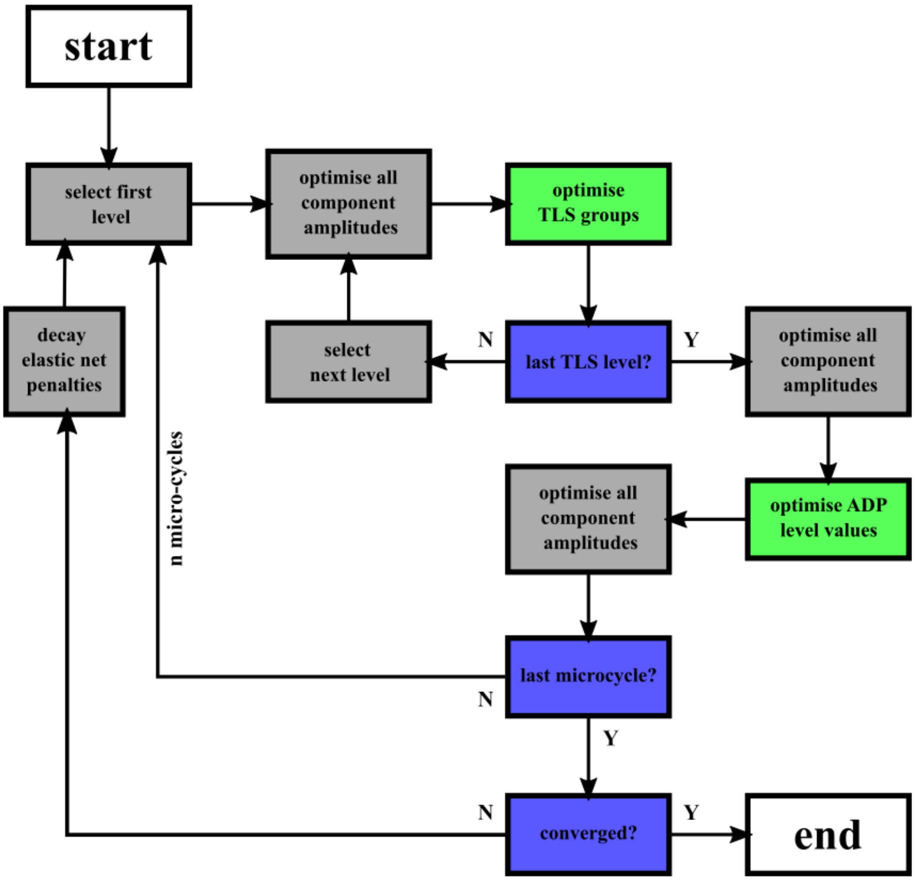


Supplementary Figure 23: **Elastic-net optimisation for echt models.**

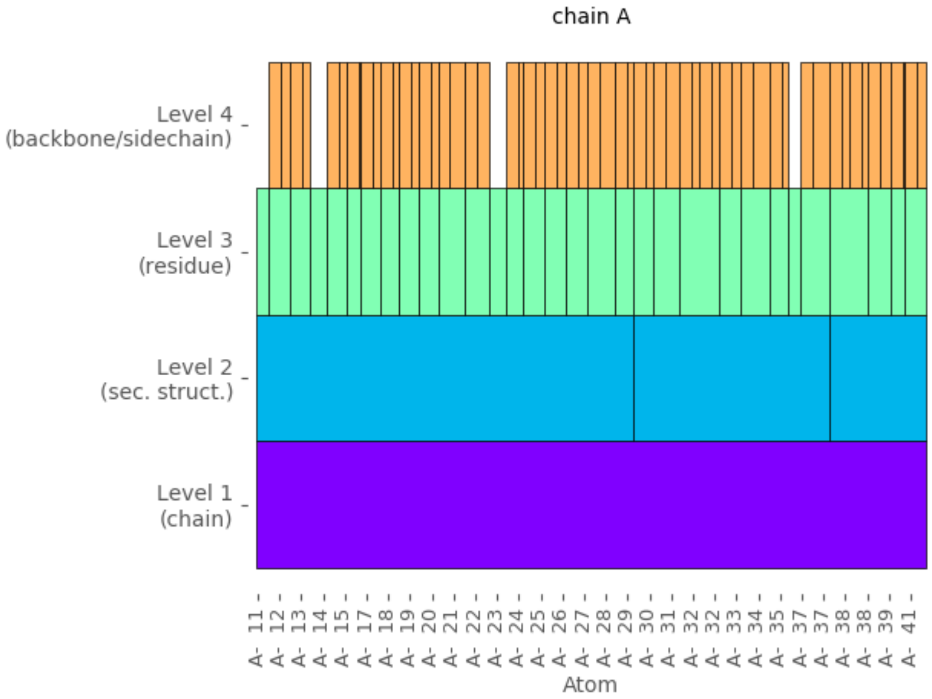


Supplementary Figure 24: **Example partitioning for residues 11-41 of 5php.** Levels are chain, secondary structure (as identified with DSSP), residue, and backbone/sidechain, respectively. Gaps in the backbone/sidechain levels are for proline or glycine or for residues with unmodelled sidechains.
